# Linked-Pair Long-Read Sequencing Strategy for Targeted Resequencing and Enrichment

**DOI:** 10.1101/2023.10.26.564243

**Authors:** Lahari Uppuluri, Christina Huan Shi, Dharma Varapula, Eleanor Young, Rachel L. Ehrlich, Yilin Wang, Danielle Piazza, Joshua Chang Mell, Kevin Y. Yip, Ming Xiao

## Abstract

In this report, we present linked-pair sequencing, a novel strategy to construct a long-read sequencing library such that adjacent fragments are linked with end-terminal duplications. We use the CRISPR-Cas9 nickase enzyme and a pool of multiple sgRNAs to perform non-random fragmentation of targeted long DNA molecules (>300kb) into smaller library-sized fragments (about 20 kbp) in a manner so as to retain physical linkage information (up to 1000 bp) between adjacent fragments. DNA molecules targeted for fragmentation are preferentially ligated with adaptors for sequencing, so this method can enrich targeted regions while taking advantage of the long-read sequencing platforms. This enables the sequencing of target regions with significantly lower total coverage, and the genome sequence within linker regions provides information for assembly and phasing. We demonstrated the validity and efficacy of the method first using phage and then by sequencing a panel of 100 full-length cancer-related genes (including both exons and introns) in the human genome. When the designed linkers contained heterozygous genetic variants, long haplotypes could be established. This sequencing strategy can be readily applied in both PacBio and Oxford Nanopore platforms. This economically viable approach is useful for targeted enrichment of hundreds of target genomic regions and where long no-gap contigs need deep sequencing.

## INTRODUCTION

The advent of high-throughput sequencing technologies has brought about significant advancement in our understanding of genomics, enabling tasks such as constructing references, identifying disease-causing variants, and uncovering both small and large-scale structural variation among genomes. While short-read sequencing stands out for its exceptional cost effectiveness and base calling accuracy, its limited read length of a few hundred base pairs (bp) presents two significant challenges in genome analysis (a) complete *de novo* genome assembly and (b) detection of large structural variations (SVs). These challenges are caused in part by the presence of large repeat families and complex genomic loci. Overcoming these challenges has led to the adoption of long-read sequencing technologies, mainly the PacBio SMRT and Oxford Nanopore technologies. One advantage of PacBio sequencing is that it provides longer reads that can cover complex and repetitive regions accurately, with an average error-corrected read length of 10-20 kbp (1). Meanwhile, Oxford Nanopore sequencing boasts an average read length of 20-50 kbp, with the ability to pass through DNA molecules as long as 1 Mbp (2,3). However, nanopore sequencing may suffer from higher error rates (2,4,5) and super-long read events are quite rare due to relatively inefficient adaptor ligation and translocation of ultra-long DNA molecules (2). Both technologies offer substantially greater sequence contiguity than short-read sequencing – longer reads span across many of the complex repetitive regions – therefore genome assembly, haplotype phasing, and calling simple and complex structural variants are vastly improved.

To generate a complete *de novo* assembly of the whole human genome, a combination of different sequencing technologies has been proven successful (6). Similarly, combining technologies has enabled the comprehensive discovery of structural variants, especially those larger than 50 bp (7). However, using multiple technology platforms to comprehensively survey the entire human genome raises project complexity and computational costs, making it impractical for broad usage, especially in clinical diagnostics. For the majority of resequencing projects, targeted approaches can be more efficient and cost-effective compared to whole genome sequencing (WGS). It is important for targeted approaches to possess a high level of multiplicity, high accuracy, and targeting flexibility across the human genome including complex regions, have straightforward workflows, and incur a low cost. Combining the benefits of targeted sequencing with long-read technologies can potentially provide ways to address both the *de novo* assembly of complex regions and the discovery of large, complex SVs. Several target enrichment approaches have been employed alongside long-read technologies, including PCR-based (8,9), computational screening (10), and commercial Cas9-based enrichment products (11). CRISPR-Cas9-mediated targeted sequencing applications are gaining popularity due to the lack of an amplification requirement and the programmability of fragmentation sites to release long-read compatible DNA fragments (12). In the same vein, nanopore sequencing is widely used for its low acquisition cost, straightforward library preparation, and for its long-read lengths.

In one early report, Cas9 was used to digest genomic DNA (2-35 µg) at locations flanking a 200 kbp target region followed by size selection of the target via pulse field gel electrophoresis. Termed CATCH, the selectively enriched target DNA (<2 ng) was long-range amplified, purified (20 µg), and 1 µg of the product was linearized, amplified for a second time, and purified, at the end of which 20-40 kbp amplicons were size-selected by pulsed-field gel electrophoresis (PFGE) before sequencing library preparation (13). Shortly after, an alternative method was reported, dubbed FLASH, that performed Cas9 cleavage on dephosphorylated bacterial genomic DNA (25-100 ng), followed by purification, dA-tailing, sequencing adaptor ligation, repurification, and amplification before Illumina sequencing (14). The number of sgRNAs used in FLASH was substantially higher than in any prior enrichment reports (in the order of 1000s), making the assay capable of detecting a large multiplicity of targets. Although this method was applied to short-read sequencing of bacterial DNA, the simplicity of the process, achieved by eliminating multiple amplification and purification steps, is worth noting. Another notable observation is the relatively low amount of starting DNA needed for this method, compared to the starting DNA necessary to physically detect, excise, and purify from PFGE. However, it is important to mention that long-read sequencing, in general, demands higher loading DNA amounts compared to short-read sequencing.

A similar method of digesting dephosphorylated genomic DNA (3 µg) with Cas9, followed by adaptor ligation was recently combined with nanopore sequencing (15). Called nCATS, this method used no PCR amplification and presented a straightforward approach for targeted nanopore sequencing. Commercial Cas9-enrichment kits for long-read sequencing appear to closely resemble this procedure. Although well-characterized, the level of multiplexing possible (as in FLASH) was not demonstrated. One of the obstacles in testing for high target multiplicity is the lack of an inexpensive and facile means to produce a large number of sgRNAs required for a complex fragmentation reaction.

Although the novel methods described above were able to selectively enrich specific genomic intervals for long-read sequencing, the Cas9/sgRNA cleavage steps eliminated the linkage between adjacent fragments, thus losing information that could be used for genome assembly or haplotype phasing across adjacent fragments. Here, we present a linked-pair sequencing strategy that is capable of sequencing long genomic regions in an enriched fashion together with greater multiplicity in targeting. Altogether, it can assemble critical genetic loci more accurately. We used the CRISPR-Cas9 editing system and carefully designed gRNA pairs to target genetic loci for enrichment. The genetic loci was tiled across by fragments matching average read lengths of the current long-read technologies for efficient sequencing. These fragments were generated by non-random, directed activity of CRISPR-Cas9 nickase to sites on opposite strands within 1 kbp and using these nicks for new DNA replication resulting in the fragments sharing tandemly duplicated terminal ends.

The linked-pair sequencing strategy not only allows the orderly fragmentation of long targeted regions into smaller library-sized fragments while retaining physical linkage information between adjacent fragments, but also enables preferential ligation of targeted DNA fragments with adaptors for sequencing for significant enrichments of targeted regions, all in an economical assay. We demonstrate the validity and efficacy of our method by first sequencing full length of phage lambda and then a panel of about 100 full-length cancer genes (including introns) in the human genome. When the designed linker sequences contained heterozygous genetic variants, long haplotypes could be established. These whole-gene haplotype sequences enable the study of genetic variants at non-coding regulatory elements and the detection of any allele-specific effects.

## RESULTS

### The principle of linked-pair sequencing

The approach relies on generating library-sized DNA fragments from long DNA molecules such that the 300-1000 bp at the ends of the adjacent DNA fragments are duplicated. An overview of the linked-pair principle is shown in Figure 1A; a long contiguous DNA molecule was non-randomly fragmented into many smaller fragments in such a way that the ends of the fragments (circled) shared the specific identical sequences up to 1000 bp (blue and green rectangles), called linkers or linker sequences. The linker sequence on the right end of fragment #1 is identical to the linker sequence on the left end of fragment 2. Fragments 1 and 2 share a linker indicated in the blue rectangle. Similarly, the right end of fragment #2 is identical to left end of fragment #3, indicated by green rectangle and so forth. When such fragments are sequenced, the overlapping linker sequences in the reads allow for the identification of their adjacent fragments. This orderly fragmentation and linker generation occurs before sequencing library preparation, and therefore the resulting DNA fragments can be processed and sequenced on current platforms, including Illumina, PacBio, and Oxford Nanopore. By implementing this method upstream of sequencing, it is possible to target and sequence many loci in the human genome, analyze long structural variations including copy numbers, and in principle resolve their haplotypes. By preserving linkers, very long genes can also be sequenced and efficiently *de novo* assembled.

**Figure 1.**
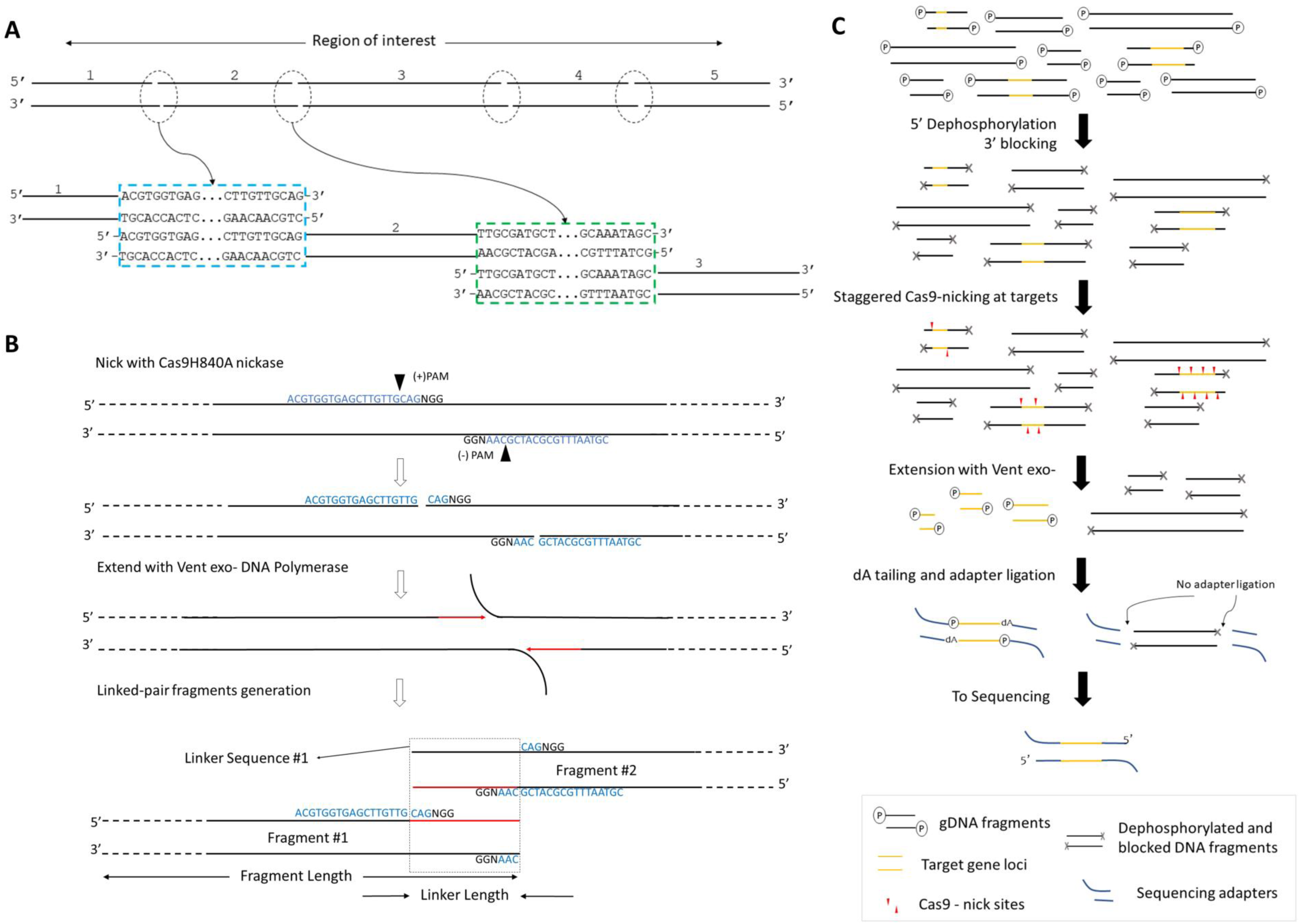
(A) Schematic showing the fundamental concept of linked pair sequencing strategy. A long DNA molecule (black bars) is deliberately fragmented in a way such that neighboring fragment pairs share end-terminal duplications. In this case, five fragments are generated. Fragments 1&2 are shown to share identical terminal sequence (linker) highlighted in blue rectangle. Fragments 2&3 are shown to share identical terminal sequence (linker) in green rectangle, and so forth. (B) Linker generation overview. Staggered nicks (black triangles) are first introduced into long DNA molecules (black bars) at specific sites (blue sequence+NGG) by Cas9 nickases. Next, strand displacing Vent exo-DNA polymerase starts new strand synthesis, which is a replication of the older strand, (red arrow) from the nick sites while translating the nicks and displacing the original strand (black-flap). The translated nicks move towards each other with new strand synthesis ultimately breaking the DNA; polymerases keep filling new strands until the end is blunt. Thus, the identical terminal sequences become linkers (identified by the dashed rectangle) connecting both fragments. (C) The workflow of constructing the linked-pair sequencing library. Input DNA (black bars) with target regions (yellow bars) are first preprocessed by 5’-dephosphorylated and 3’-blocking. Preprocessed samples without a phosphate group at the 5’ end and without a hydroxyl group at the 3’ end (gray cross-black bars) are nicked with Cas9-nickase-sgRNA complexes to introduce staggered nicks (red triangles) at target loci. Vent exo-is used to generate linked-pair fragments of the target loci. These fragments are subject to regular library construction protocol, dA-tailing and ligation with sequencing-adapters (blue bars, Y-shaped). Once ligated, these fragments can be sequenced on a flow cell. Sequencing adapters will not be attached to non-target fragments.

An overview of linker-pair fragmentation is shown in Figure 1-B. First, a pair of staggered nicks on opposing strands (black triangles arrows in Figure 1B) in regions of interest on high molecular weight DNA molecules. These nicks are up to1000 bp apart and made using the mutant Cas9D10A or Cas9H840A protein with two unique sgRNAs. One such nick pair by Cas9 H840A nickase is shown in Figure 1B. Then, DNA polymerases with strand displacing capabilities like Klenow (exo-) or Vent (exo-) polymerase are used to synthesize new DNA strands from both nick sites while displacing the original strands. New synthesis, indicated as red arrows, is replicated off the original strands. As these two newly synthesized strands extend toward each other, dsDNA breaks off while DNA polymerase continues to fill in the single-stranded 5’ overhangs (black flaps in Figure 1B). Consequently, the two fragments share identical tandem duplicated sequence, highlighted by a rectangle at the fragment ends in Figure 1B and a linker is generated by a pair of sgRNA nick sites. Two pairs of nick sites together generate a linked-pair fragment. Length of linker is defined as the distance between NGG −3 bases of the upstream nick site to NGG +3 bases of the downstream nick site within the same linker region. Outermost fragments at each locus of interest receives only one linker. Fragment length is defined as distance between two linkers (linkers included). In case of single-linker fragments, the length becomes distance to nearest end (Figure 1B).

Schematic in Figure 1C shows the complete workflow for a typical linked-pair library preparation of more complex genome like bacterial or human DNA. The process starts with minimally fragmented high molecular weight DNA (black bars) sample for input. Sample is pre-processed with 5’ dephosphorylation and 3’ blocking of randomly available ends in DNA. Dephosphorylation with a phosphatase enzyme removes a phosphate group from all available 5’ ends (indicated as (P) in Figure 1C). 3’ ends are rendered inactive by incorporation of dideoxynucleotides by Klenow exo-DNA polymerase. These preprocessing steps (indicated by grey cross on DNA) are crucial to prevent sequencing-adaptor ligation to non-target DNA and minimizing sequencing of off-target DNA fragments. Next, staggered nick site pairs (red triangles) are introduced into loci of interest (yellow stretches) on pre-processed long DNA fragments by an appropriate Cas9 nickase. Vent (exo-) polymerase is used to extend from nick sites towards each other resulting in controlled, non-random fragmentation of target regions into library-sized fragments. These fragments are cleaned up and dA-tailed before ligation. Freshly fragmented DNA retains active 5’ and 3’ ends which are available and are preferably tagged with a sequencing-adapter. Thus, prepared library is put on a sequencer where fragments which have an adapter tagged can be sequenced.

### Performance of linked-pair sequencing in model systems

We first used Lambda phage DNA (48.5 kbp) as model systems to demonstrate linked-pair sequencing and characterize two key parameters, fragmentation efficiency and contiguity without gaps. A set of six sgRNA pairs was designed to cover the linear lambda DNA, generating seven linked-pair fragments, i.e., fragments connected by overlapping linkers, ranging from 1 kbp to 13 kbp connected by linkers ranging from 50 bp to 250 bp (Table 1, Supplementary Table 1). Figure 2A shows the schematic design of the sgRNAs. The nick sites generated by 6 pairs of gRNAs are shown as black arrows and the six linker sequences are depicted as yellow dots on the Lambda DNA (gray bar). The linked pair fragments then were generated by first nicking 1000 ng of Lambda DNA with Cas9D10A nickase and then extending with Vent (exo-) DNA polymerase. The generated product was electrophoresed on 1% agarose. The gel image in Figure 2B confirmed that correctly sized fragments were generated.

**Table 1.** Fragmentation efficiency of Lambda DNA with linked-pair strategy.

| First sgRNA at linker site (Location bp) | First sgRNA at linker site (Location bp) | Linker Length(bp) | Fragment Length (bp) | reads with Linkers | reads without Linkers | Fragmentation Efficiency |
| --- | --- | --- | --- | --- | --- | --- |
| Expt. 1 |  |  |  |  |  |  |
| sgRNA1(6270) | sgRNA2(6359) | 83 | 6359 | 4013 | 526 | 88.4% |
| sgRNA3(13090) | sgRNA4(13164) | 68 | 6894 | 2473 | 935 | 64.2% |
| sgRNA5(24569) | sgRNA6(24649) | 74 | 11559 | 2161 | 238 | 65.4% |
| sgRNA7(27154) | sgRNA8(27431) | 81 | 2862 | 6440 | 2053 | 68.3% |
| sgRNA9(34261) | sgRNA10(34495) | 231 | 7341 | 2591 | 3511 | 32.2% |
| sgRNA11(47446) | sgRNA12(47586) | 134 | 13325 | 2197 | 531 | 34.2% |
|  |  |  | 1056 | 8576 |  | 80.5% |
| Expt. 2 |  |  |  |  |  |  |
| sgRNA13(24629) | sgRNA14(24728) | 99 |  | 330 | 232 |  |
| sgRNA15(26560) | sgRNA16(27160) | 600 | 2531 | 1195 | 20 | 98.4% |
| Expt. 3 |  |  |  |  |  |  |
| sgRNA17(21980) | sgRNA18(23010) | 1030 |  | 27 | 42 |  |
| sgRNA19(24629) | sgRNA20(24725) | 96 | 2745 | 629 | 259 | 70.8% |

**Figure 2.**
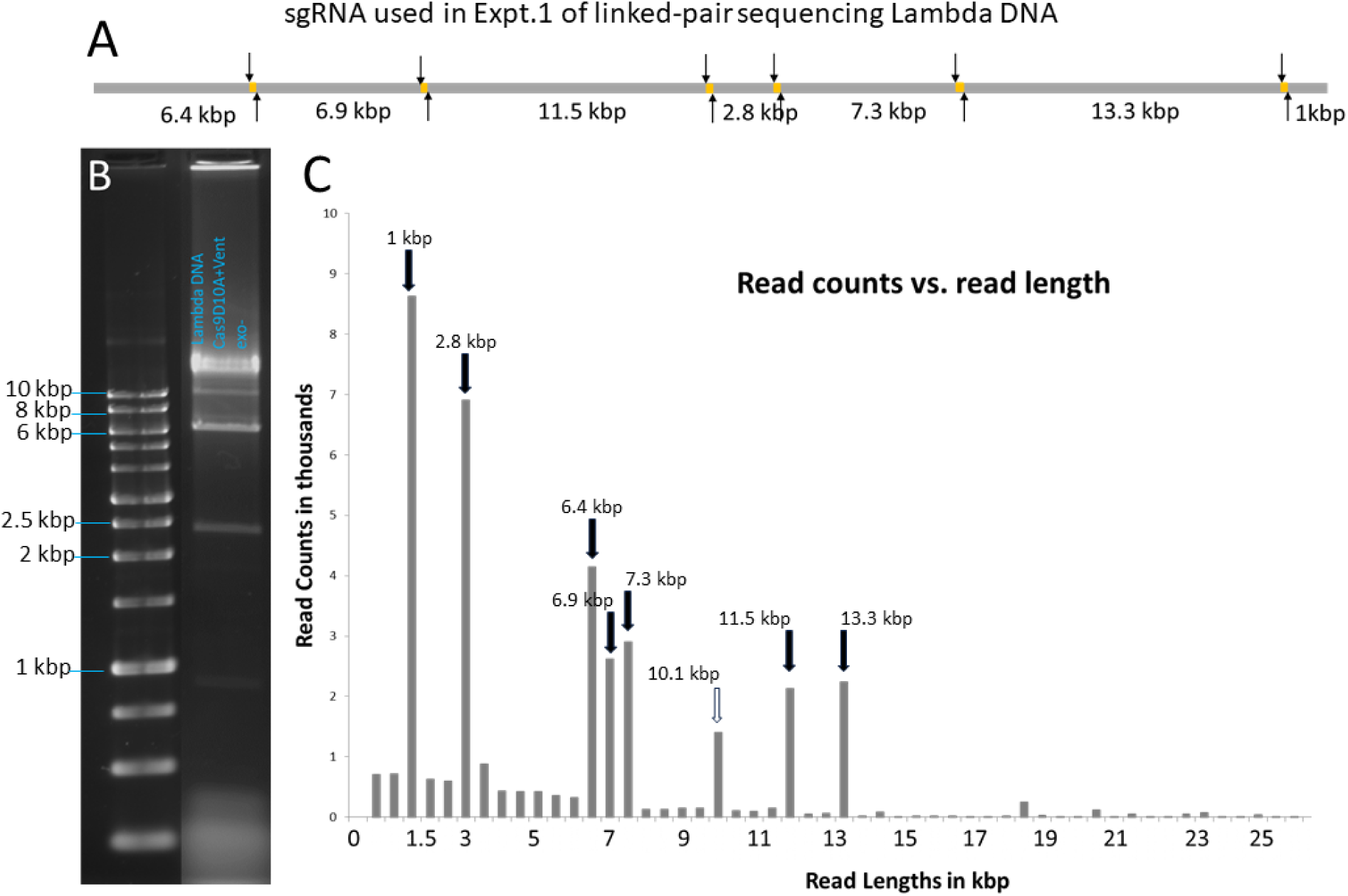
Lambda genome was fragmented and sequenced following the linked-pair strategy. (A) shows a schematic with the linker sequences (yellow dots) anticipated by the designed six sgRNA pairs (arrows) along Lambda DNA backbone (Gray line). It will generate 7 fragments with lengths varying between 2.8kb and 11.5kb. (B) Agarose gel images. The Lambda linked-pair fragments produced are a good match with the design shown in Figure 2A. (C) Oxford nanopore sequencing results of Lambda linked-pair library. It shows a histogram of sequencing read lengths from this experiment. X-axis represents read lengths and Y-axis represents read counts. Dark arrows show peaks for linked-pair fragments corresponding to anticipated linked-pair fragments. Unexpected 10.1 kb (white arrow) at peak is from unfragmented linker between 2.8 kb and 7.3 kb fragments.

Fragments of 1, 2.5, 7, 11.5, and 13.3 kbp were observed as discrete bands, whereas the 6.4 and 6.9 kbp fragments were not clearly resolved and resulted in a merged band of higher intensity. The peak at 10 kbp could be arising from non-fragmented reads at the 4^th^ linker. The remaining sample was cleaned before adapter ligation and loading on the nanopore flow cell. In Figure 2C, a histogram of sequencing read lengths from this experiment is shown, in which read counts correspond to those that fully cover an anticipated fragment. All peaks from the histogram mapped to the anticipated fragment lengths from the sgRNA design.

A snapshot of a subset of aligned reads with linkers is shown in Figure 3. Panel 3A shows sgRNA pairs with nick sites, linker sequences, and anticipated fragments indicated. Panel 3B has the coverage track followed by a subset of all sequenced reads. On analyzing the alignment, an increase in coverage at all six overlapping linkers (yellow dots) was observed, due to the generation of duplicated sequences during non-random fragmentation. Two examples of linkers can be observed in Figure 3C, which shows a zoomed-in view of the sequencing reads. For linker 1, the top set of reads can be seen extending from the left and terminating with the linker, whereas the bottom set of reads begins with the linker and extends to the right showing successful generation of linked pair reads.

**Figure 3.**
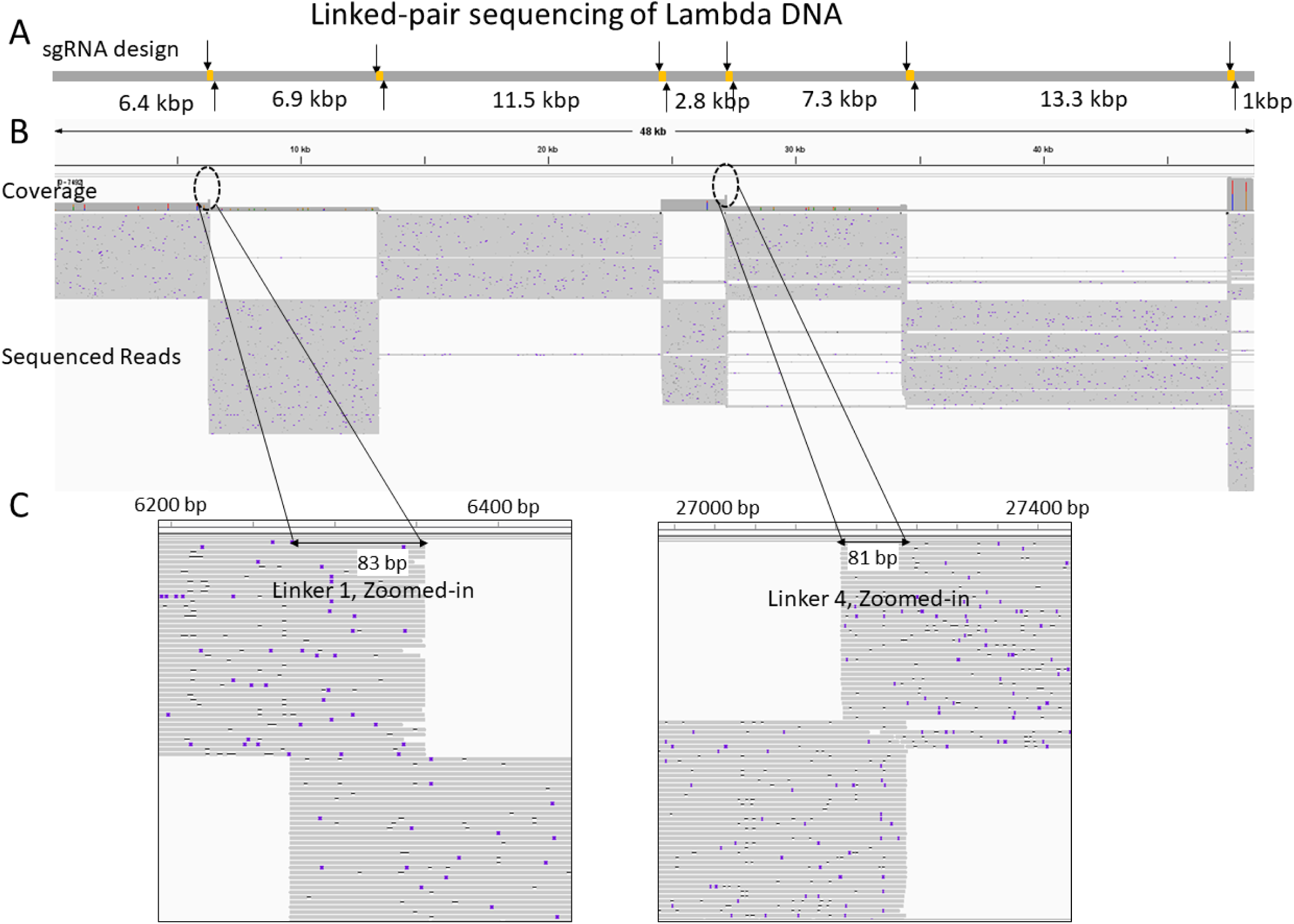
Visualization of sequencing reads of Lambda linked pair fragments in Expt. 1. Panel A shows designed sgRNA (arrows) and anticipated linker sequences (yellow dots) on the Lambda reference genome (gray bar). Panel B shows a track with the depth of reads coverage along Lambda DNA reference followed by a subset of sequencing reads (grey lines) aligned to the Lambda reference. Panel C shows zoomed-in regions (circles in panel B) where the neighboring fragments have identical linker sequences. Linker sequence #1 (left) is shared between fragments 6.4 kb and 6.9 kb (panel A) and linker sequence #4 is (right) is shared between fragments 2.8 kb and 7.3 kb (panel A).

To assess the system performance, the percentage of reads containing the linker sequences between any two pairs of nick sites, i.e., fragmentation efficiency, was determined and reported in Table 1. For calculating the fragmentation efficiencies, we accounted for the sequenced reads arising from DNA molecules whose ends mapped to predicted linkers or instead mapped across the linkers with no evidence of fragmentation or duplication.

For example, we can calculate the fragmentation efficiency of the first linker by counting reads. Between the 1st base of Lambda DNA and the first linker (sgRNA1 at 6270 bp and sgRNA2 at 6359 bp), there were 4013 reads with linkers and a read length of ∼6.3 kbp. Also, 526 reads were mapped across this linker without any break. Thus, the fragmentation efficiency at this site was computed to be 4013/ (4013+526) =88.41% (a fraction of the reads with linkers among the total sequenced reads). Similarly, the fragmentation efficiency of the 6.9 kbp fragment between the first pair of nicks and the second pair of nicks (sgRNA3 and sgRNA4) was observed to be 64.16% after accounting for linker generation on both ends of the fragment. While an average fragmentation efficiency of 62.1% was observed, fragments flanking the 231 bp linker had lower efficiency. However, reads with the correct end-terminal linkers adjacent to this 231 bp linker were high at 76% and 81% respectively. We supposed that this decrease could either have been due to the on-target nicking efficiencies of sgRNA 9 or sgRNA10. An inefficient extension of the polymerase in extending to a full linker length of 231 bp and ultimately fragmenting the genome was also a possibility.

We then designed additional pairs of sgRNAs to evaluate the feasibility and efficiency of generating longer linkers. The sgRNA13 – sgRNA16 pairs generated one 600 bp linker and one 99 bp linker on Lambda DNA, and sgRNA17– sgRNA20 generated one 1 kbp linker and one 96 bp linker. Independent experiments (Expt. 2 and Expt. 3 in Table 1) were conducted, showing that the fragmentation efficiency for these two longer linkers was as high as that of the smaller linkers. This suggests that the lower fragmentation efficiency of the 231 bp linker for Expt. 1 was likely due to the lower on-target nicking efficiencies of sgRNA 9 or sgRNA10designed for this linker.

### Enrichment of full-length cancer gene panel with linked-pair targeted sequencing

In our initial experiments with Lambda model system, our protocol was successful in performing linked-pair sequencing with linkers up to 1000 bp. To scale up as well as to characterize the on-target enrichment of our linked-pair sequencing protocol, we attempted to develop linked-pair protocols to perform targeted sequencing in the human genome. A set of 103 cancer genes were shortlisted from a panel of frequently mutated cancer genes (16,17), and 384 sgRNA sequences (Supplementary Tables 2 and 3) were designed to fragment and sequence DNA using the linked-pair method with Oxford Nanopore sequencing. Since the human genome is several thousand-fold larger than our model systems in addition to having several stretches of complex and hard-to-sequence regions, there will be an increased likelihood of sequencing non-target fragments. To suppress the availability of non-target fragments to sequencing adapter ligation, we first treated the DNA samples with a 5’ dephosphorylating enzyme to dephosphorylate any randomly available 5’ ends due to internal nicks or damage. (15) To further suppress the background and improve reaction efficiency, we introduced dideoxynucleotides at all available 3’-ends to inhibit non-specific extensions and therefore ligations. Adding these pre-processing steps to our earlier protocol increased target enrichment (Figure 1C) by increasing the likelihood of selective ligation of adapters to target fragments which are favorably sized to translate through nanopores. We also assayed and optimized other parameters including reaction buffers, sgRNA, and enzyme concentrations to improve the linked-pair protocol for the human genome. The same is also reported in the methods section.

We developed a computational method for designing sgRNAs that produce DNA fragments covering specific target genomic loci (detailed in methods section). Briefly, for each target locus too long to be covered by a single sequencing read, neighboring DNA fragments shared a common linker such that they could be easily assembled. Our method considered various factors in the design, including read length, linker length, off-target effects of sgRNA targeting, and genetic polymorphisms that may influence on-target binding or facilitate haplotype reconstruction. This method was applied to design 384 sgRNAs to produce a total of 160 DNA fragments that cover 103 cancer genes (Supplementary Tables 2 and 3). The panel included 29 long genes which were designed to be covered by two or more reads and 74 shorter genes were designed to be covered by a single read. The designed 384 gRNAs together were expected to produce a total of 160 fragments of ∼15 kbp length. For genes covered by more than one fragment, the fragments were designed to be connected by linkers 200 – 500bp in size.

Based on our design, the linked-pair sequencing library was generated in two sequential stages from the human lymphoblastoid cell line NA12878, i.e., by sequential fragmentation of the short genes followed by the long and short genes. A total of 1,083,376 reads with an average read length of 12,066 bp were produced resulting in a total of 13,071,923,552 bp after adapter trimming, corresponding to a 4.2x average genome-wide coverage. To understand the origin of these reads, we aligned them all to the reference human genome GRCh38. About 99.1% of reads produced aligned to the reference. The aligned sequencing reads fully covered all the designed linker regions and target genes, resulting in on-target coverage of 100%. Next, we studied the enrichment of sequencing reads at the target cancer gene loci. The average depth of coverage at the gene body region (from the first transcription start site [TSS] to the last transcription termination site [TTS]) was 30.9x (Figure 4A), which was seven-fold higher than the baseline genome-wide coverage. In our gRNA design, to fully cover the cancer gene set, the terminal linkers were set outside the gene body regions. As a result, some flanking regions were also enriched, and depth-of-coverage gradually decreased to the genome-wide average as a function of distance from the gene body region (Figure 4A). When we focused on whole target regions, i.e., the genomic spans covered by the expected DNA fragments, rather than the gene body regions only, we got an even bigger contrast of the depth of coverage between the target regions and the flanking regions (Figure 4B).

**Figure 4.**
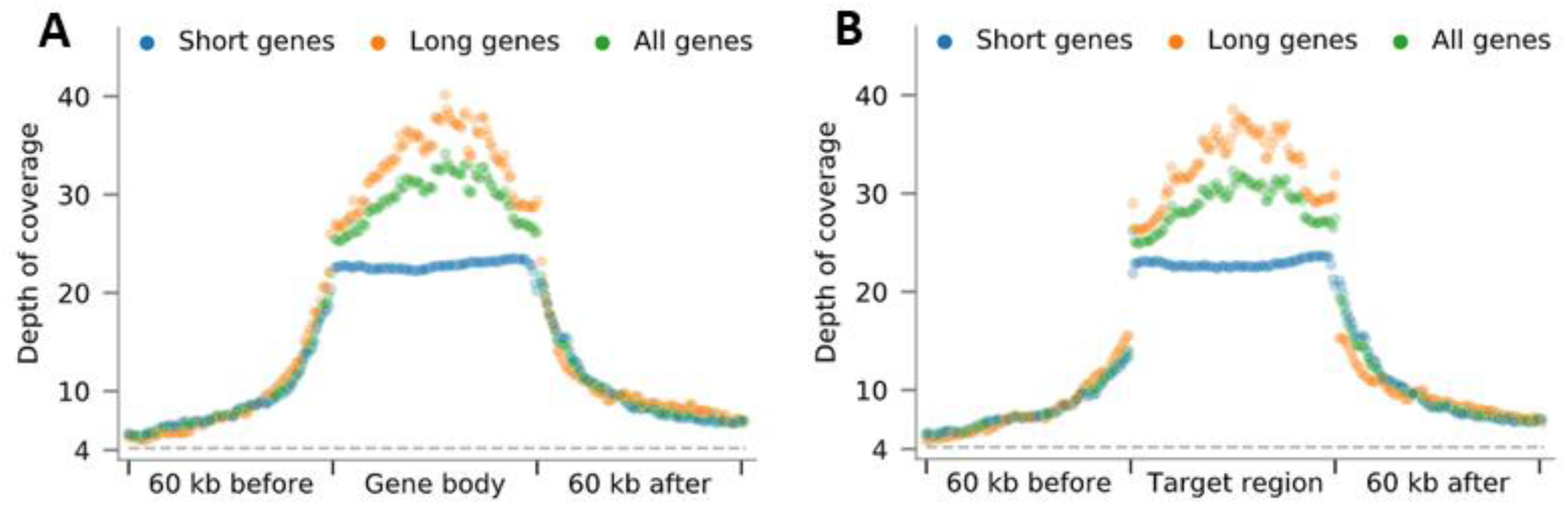
(A-B) Enrichment of aligned sequencing reads at all the cancer gene loci included in our design. (A) Each gene body region (from TSS to TTS) or (B) target region (from first sgRNA to last sgRNA) and each 20kb flanking region was divided into 100 equal-sized sub-regions and the average depth of coverage of each sub-region, over all included genes, only the short genes, or only the long genes, is shown as a single dot.

An example of reads aligned to one of the long genes, *NBN* (Chr8: 89,933,335-89,984,724) as viewed in IGV is shown in Figure 5. The gene locus is annotated at the top (gray bar) and has >50x coverage. The pairs of gRNAs, shown as black arrows, indicated the nick sites for generating the four linkers (yellow dots). The black arrows correspond to the spikes in coverage at four locations as a result of overlapping linkers. Two dashed circles shown in the coverage track indicate amplified coverage due to the linker sequences of both fragments. The bottom two panels of Figure 5 show a zoomed-in view of the 2^nd^ and the 3^rd^ linker sequences on the *NBN* gene. In each zoomed-in panel, the top fragments can be seen extending from the left and terminating with the linker while the bottom fragments begin with the linker and extend to the right. In this experiment, the non-target region has an average of ∼4x sequencing depth, while the coverage of the targeted region has much higher average coverage resulting in 7x enrichment. Altogether, our results demonstrate the proof-in principle for targeted sequencing using linked-pair sequencing strategy.

**Figure 5.**
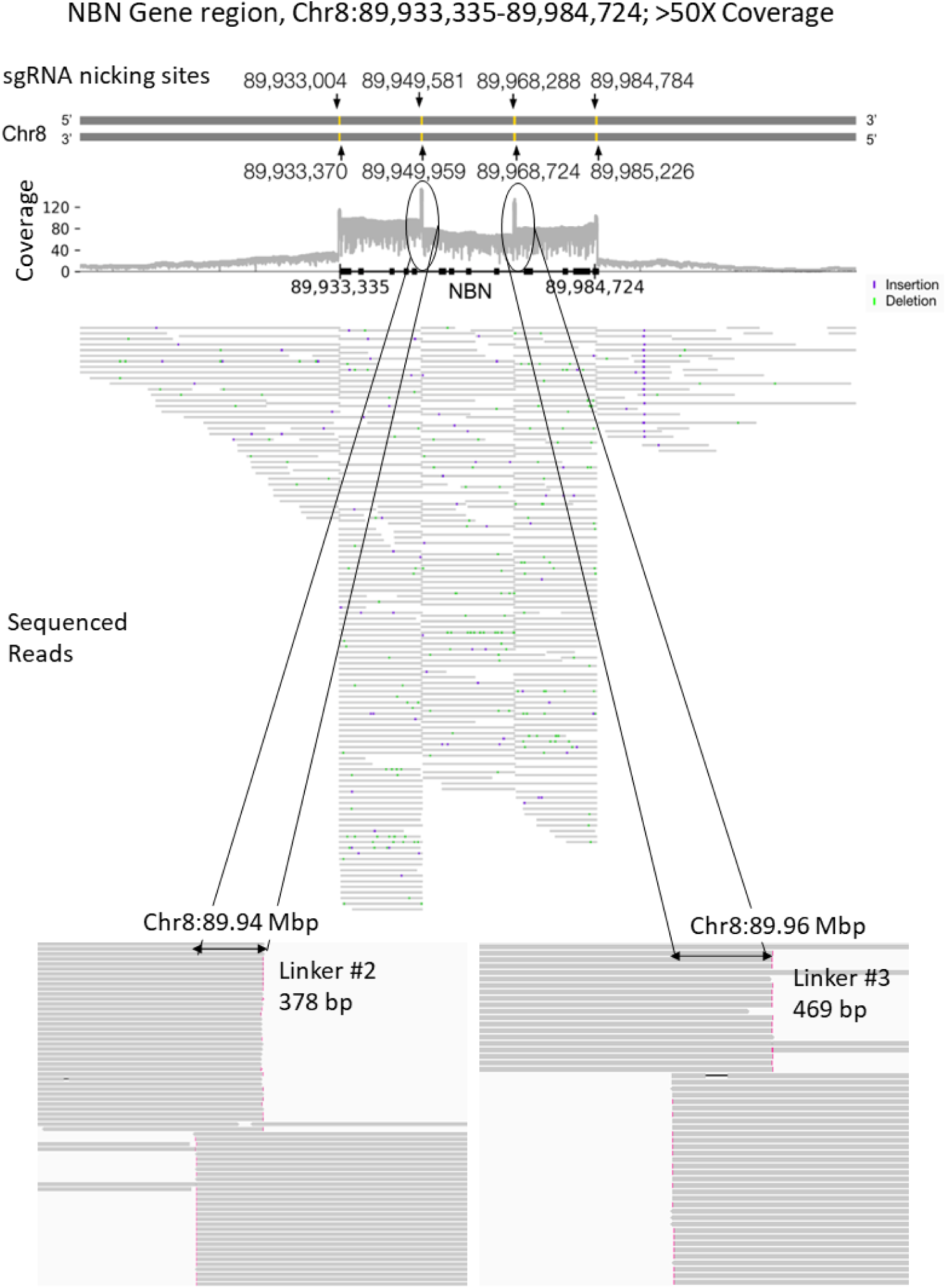
Sequencing and read alignment of the NBN cancer gene (double-headed arrow) on chromosome 8 with our linked-pair strategy. The top two lines show that four linkers (yellow bars) are created by four corresponding pairs of gRNAs (black arrows). The coverage plot below them shows the number of aligned reads covering each genomic position, with clear increase of read coverage at the linker regions observed. The next part shows alignments of individual reads (gray horizontal lines) to the NBN locus, which cover both the exons and introns. The last part shows zoomed-in views of the ends of reads aligned near the second and third linkers, which clearly show the overlap of adjacent DNA fragments due to the linkers.

Next, we used the read alignments to reconstruct the haplotypes of two long genes, *NBN* (Figure 6) and *PIK3R1* (Supplementary Figure 1). For the *NBN* gene, we designed 4 pairs of sgRNAs that would cover the whole gene with 5 DNA fragments, where each adjacent pair of fragments shared a linker. The actual sequencing reads produced were indeed enriched in the gene body region and there are clear peaks of increased coverage at the linker regions (Figure 6A). Within the linker regions, we identified 6 heterozygous single-nucleotide variants (SNVs) by comparing our reads with the reference and phased them into two haplotypes found in the GM12878 reference genome (Figure 6B-C) (18). We used a graph-based method to reconstruct the haplotypes using our sequencing reads detailed in methods section). Briefly, for any two SNV sites, if they were covered by the same sequencing read, we put an edge between their alleles whose weight was equal to the number of reads with both alleles. These edges together formed a graph (Figure 6E). By selecting only allele pairs with a statistically significant number of supporting reads (p<0.05, Fisher’s exact test), two separate haplotypes are formed, which are consistent with the haplotypes determined by GIAB (Figure 6E). For the *PIK3R1* gene, we designed 7 pairs of gRNAs that would cover the whole gene with 8 DNA fragments, again with every pair of adjacent fragments sharing a linker (Supplementary Figure 2A). Among these 7 linkers, the first and last ones are outside the gene body region, while the remaining 5 are inside it (Supplementary Figure 1B). Within these 5 linkers, we identified 4 heterozygous SNVs and 1 heterozygous deletion, which had also been identified by the GIAB project. Using the same graph-based approach, we phased these 5 variants into two haplotypes, which were consistent with the GIAB haplotypes (Supplementary Figure 1C-E).

**Figure 6.**
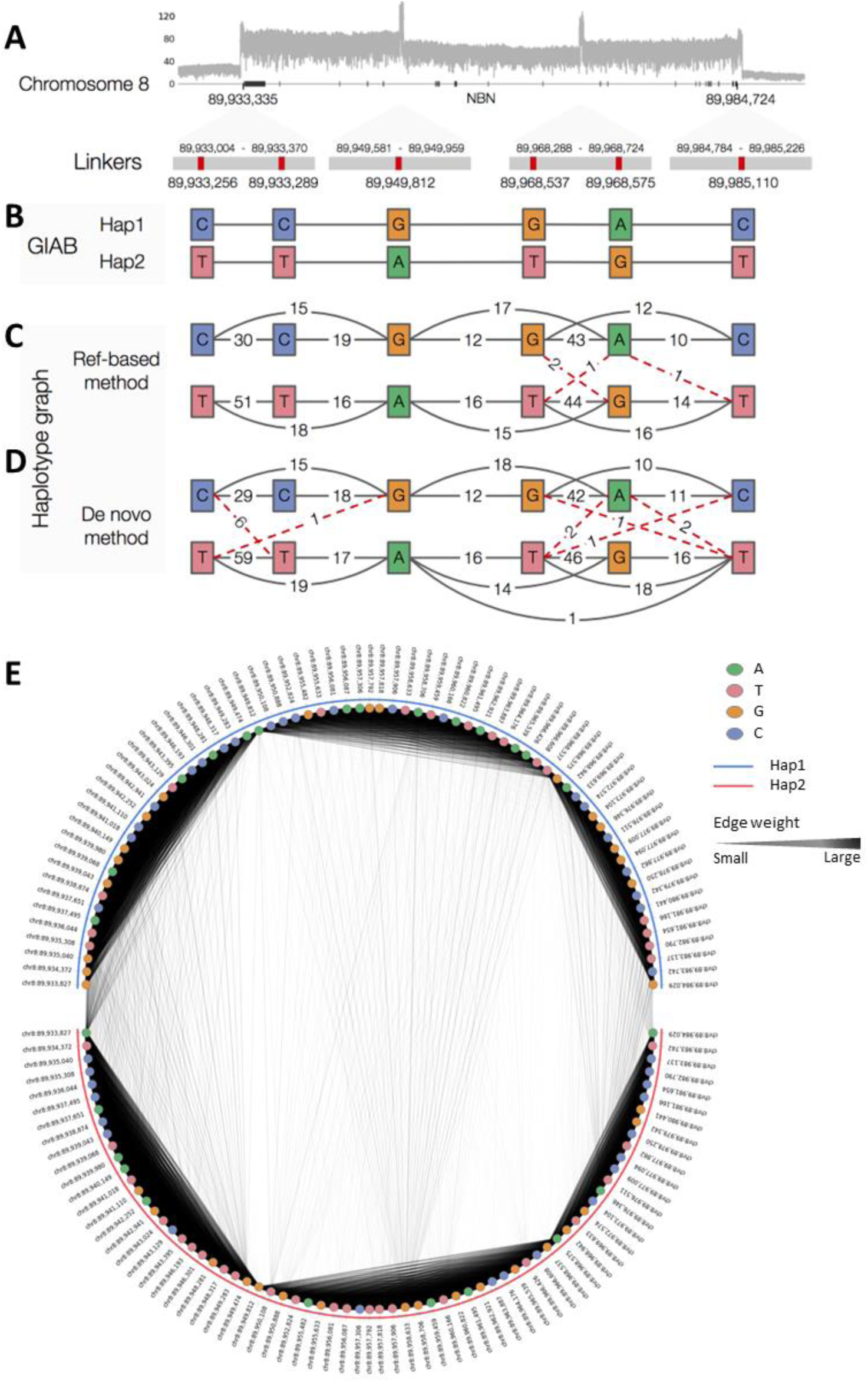
Haplotype phasing of the heterozygous variants within the linker regions of the gene NBN. (A) Depth of sequencing reads aligned to the gene region. (B) The six heterozygous SNVs within the four linkers and the corresponding haplotypes phased by GIAB. (C-D) Haplotype graphs constructed (C) by aligning the sequencing reads to the reference genome and (D) by multiple sequence alignment of the sequencing reads. Each solid black line indicates two alleles found on the same read that are from the same GIAB haplotype, and each dotted red line indicates two alleles found on the same read that are from different GIAB haplotypes. Each of these two types of lines is labeled by the number of supporting reads. (E) Haplotype phasing of all heterozygous SNVs within the genes. In each panel, the variant sites are ordered from left to right according to their genomic locations. The two alleles from each variant site are placed at either the upper or lower semi-circle, according to the haplotype that it was assigned to by our method. Both the width and darkness of each edge reflect the number of reads in which the two alleles that define the edge co-appear.

We further explored the possibility of *de novo* haplotype reconstruction without making use of a reference genome. The main idea is still to connect variant alleles that appear in different linkers of a sequencing read, but instead of defining the variant sites by aligning sequencing reads to a reference, we defined them by performing multiple sequence alignments of the sequencing reads themselves (detailed in methods section). For the *NBN* and *PIK3R1* genes, the resulting haplotypes were consistent with the ones from GIAB (Figure 6D and Supplementary Figure 1D).

For this design, we considered only heterozygous variants within the linkers, which were specifically designed by our algorithm to serve as anchor points for phasing the haplotypes of long genes. On the other hand, genes can also contain heterozygous variants outside the linkers. To see if our linked sequencing method can also phase the haplotypes of these variants, we identified all heterozygous SNVs within the gene body regions of *NBN* and *PIK3R1* from the read alignments. Again, we connected alleles at different variant sites by an edge whose weight was the number of reads in which the alleles cooccurred. For the *NBN* gene, we identified 71 heterozygous SNVs from its gene body, and successfully phased 69 of them into two haplotypes that were fully consistent with GIAB haplotypes (Figure 6A). The remaining 2 SNVs were not identified by GIAB. For the *PIK3R1* gene, we identified 61 heterozygous SNVs from its gene body. We phased 59 of them into two haplotypes, of which 58 were phased in the same way as GIAB (Supplementary Figure 1D). The remaining SNV was not identified by GIAB.

## DISCUSSION

The current long-read platforms can obtain sequencing reads upwards of 100 kbp - 1Mbp+, but it is important to note that the frequency of these reads is significantly lower than that of ∼20 kbp reads, which is closer to the average read length of both Nanopore and Pacbio platforms (2). Shorter molecules are favored for getting trapped by a sequencing pore or a SMRT well resulting in a trade-off between read lengths and throughput. As a result, longer molecules in a library contribute to lower sequencing yield. Targeted sequencing on top of this restricts the available number of molecules resulting in wasted throughput. Several groups have overcome this by using CRISPR-Cas based target enrichment methods due to their programmable nature and molecule-length-independence for targeting unlike hybridization-based capture or PCR enrichment (12). There is continued interest in enrichment methods that are economical facile.

We report here a linked-pair sequencing strategy, a new approach in which the CRISPR-Cas9 editing system is leveraged to perform programmable non-random fragmentation of targets into favorably sized fragments for sequencing on long read platforms. The sequencing library constructed using these fragments maintains the contiguity of reads through the tandem duplicated sequences at fragment ends and improves the sequencing efficiency of targeted regions.

The simple lambda phage model was utilized for proof-of-concept experiments. Here, both the contiguity of alignment without gaps as well as fragmentation efficiency were measured. The sequencing library constructed using these fragments maintains the contiguity of reads through the tandem duplicated sequences at fragment ends and improves the sequencing efficiency of targeted regions. Together, this suggests the success of linked-pair fragmentation protocol in tiling across a genomic stretch.

Next, we validated our approach for targeting sequencing the entire gene region including introns, exons, upstream and downstream flanking regions of a large heterogenous cancer gene panel comprising of a mix of long and short genes. Our experiments showed that the long genes were sequenced to an average depth of 34.7x and the short genes to an average depth of 23.9x. In contrast to other enrichment methods, it is worth noting that entire throughput generated has been generated in a single experiment. With fragments matching average read length of sequencing platforms, the likelihood of capture of target regions for sequencing is higher compared ultralong molecules. Our approach enables more information about targeted genes to be obtained with little to no extra cost compared to a whole exome sequencing assay or whole genome assay without sacrificed throughput. Coupled with improved error rates of nanopore and pacbio technologies, linked-pair fragmentation could be readily applied to targeted resequencing, diagnostic gene panels to identify missed driving mutations, long range haplotyping of various targets in the microbial or human genomes (19–22).

Linkers are a unique feature of our approach. Iterative gRNA designs and experimentation with Lambda DNA indicate that long linkers up to 1000 bp could be generated with our protocol. It is crucial that the Cas9-sgRNA has efficient on-target nicking activity and that the polymerase has strong displacement activity. Both Cas9D10A and Cas9H840A worked equally well while the Vent (exo-) polymerase performed the best among the polymerases tested. On comparing the performance of sgRNA vs. cRNA/tracrRNA (23,24), we found that both performed well (data not shown). Linkers designed around variants could be used to resolve haplotypes and detect minor alleles.

Lastly, it is common practice to utilize tens of thousands of sgRNAs to expand multiplexed high throughput CRISPR-Cas9 applications (25). In vitro synthesis of these guides contributes to cost savings in our assay. We designed a total of 384 gRNAs, 119 gRNA pairs for long genes and 73 pairs for short genes. In the same vein, a few hundred up to a thousand guides has already been reported in the literature (23,26). There are several commercial companies, such as GeneScript and Twist Bioscience, that offer custom synthesis services for up to 90,000 sgRNAs. According to our preliminary analysis, this amount is sufficient for sequencing all the complete genes of the human genome, including introns and flanking intervals, with a linked-pair sequencing strategy. It is worth noting that the species purity and concentration in an sgRNA pool can significantly affect the fragmentation efficiency and thus the performance.

## METHODS

### DNA preparation

For this report, Lambda DNA was ordered from New England Biolabs (NEB). For experiments on the human genome, NA12878 cells (Coriell Institute) were taken from culture and their DNA was extracted using a Nanobind disk-based solid phase extraction kit (Bionano Genomics) as per manufacturer’s recommendations. The DNA samples were quantified before fragmentation reactions using Qubit and an AccuGreen Broad Range dsDNA Quantitation Kit (Biotium).

### Single guide RNA design for Lambda experiments

For this design, pairs of gRNAs, with one gRNA nicking the positive strand and one nicking the negative strand such that the linker sequences were 50-250 bp for Lambda were selected. Another constraint was that the fragment length be under 25 kbp. The Lambda reference was scanned for all feasible 20mer-NGG sequences. The on- and off-target scores of these guides were checked on IDT’s CRISPR-Cas9 gRNA checker tool (www.idtdna.com/site/order/designtool/index/CRISPR_CUSTOM) and the guides with the best scores and most dissimilarity amongst the pool were shortlisted.

### Single guide RNA synthesis

Oligomers encoded with a T7 promoter (5’-TTCTAATACGACTCACTATAG), a 20mer single guide RNA (sgRNA) sequence, and a universal overlap sequence (5’-GTTTTAGAGCTAGA) were designed and ordered from IDT. These oligos were normalized in concentration, pooled, and hybridized to a universal 85-base oligo at the overlap, and extended to form dsDNA. These double-stranded oligos acted as templates for a subsequent *in vitro* transcription reaction in which the sgRNAs for the linked-pair sequencing were generated. Briefly, a hybridization reaction was carried out in 1X Buffer 2 (NEB). 10 µM pool of designed oligomers and 10 µM of a complementary-overlap-containing-oligomer were first denatured at 95°C for 15 seconds and allowed to hybridize at 43°C for 5min. The hybridized oligos were then extended with 5U of Klenow (exo-) at 37°C for 1h in the presence of 2mM dNTPs. Next, an exonuclease treatment was carried out at 37°C for 1h with 10U of exonuclease I (NEB) in 1X exonuclease buffer (NEB). The dsDNA was then purified using a Qiagen Nucleotide removal kit, and quality and quantity were assessed using UV-Vis spectroscopy on a Synergy H1 plate reader (Biotek). A transcription reaction was then carried out on the purified and quantified dsDNA using the T7 HiScribe transcription kit (NEB). The T7 RNA Polymerase recognizes the T7 promoter region that seeds transcription of the adjacent 20-mer target sequence thus generating the targeting sgRNA for Cas9-mediated nicking of genomic DNA templates. The synthesized sgRNAs were purified using a Monarch RNA purification kit (NEB), and quality and quantity were assessed using UV-Vis spectroscopy on a Synergy H1 plate reader (Biotek). Purified dsDNA and sgRNA were stored at −20°C and found to function for at least 3 months.

### Generation of lambda phage linked-pair fragments

First, 400ng of Cas9nickase -Cas9H840A (IDT) or Cas9D10A (NEB) was pre-incubated in 1× NEBuffer 3.1 (NEB) with 25 pmol sgRNA at 37°C for 15min to form Cas9-Ribonucleoprotein complex. Then, 1000 ng of Lambda Phage (NEB) was added to the tube, and a nicking reaction was carried out at 37°C for 2 h. Nicked DNA was then extended with 3 U of Vent (exo-) Polymerase (NEB), 100-300 µM dNTPs, and 1× Thermopol (NEB) at 72°C for 60 min. After extension, the reaction was purified twice with AMPURE XP beads and was assessed on 1% agarose gel before proceeding with sequencing library preparation.

### Gel electrophoresis

After fragmentation and before library preparation, fragmented and purified DNA samples were assessed by running electrophoresis using a 1% agarose gel slab in 1X TAE buffer at 100 V for 75 minutes. DNA was stained with 1X SYBR™ Safe stain (Invitrogen) and visualized on UVP GelStudio (Analytik Jena) in epifluorescence mode.

### Preparation of nanopore sequencing library

About 0.5-1 µg of Lambda or 3-5 µg of purified NA12878 (Coriell Institute) fragments were used as input for sequencing library preparation. A dA-tailing reaction was performed in the presence of 1mM dATPs and 1X Thermopol (NEB) with 5U of Taq DNA Polymerase 72°C for 5m. The manufacturer-recommended amounts of sequencing adapters and ligation buffer from the SQK-LSK109 kit (Oxford Nanopore) were added to this reaction along with NEBNext Quick T4DNA Ligase. The ligation reaction was carried on at room temperature for 15 minutes before AMPURE XP beads were cleaned up. The library was quantified with a Qubit fluorometer before mixing with sequencing buffer loading beads and loading on a primed flow cell. Priming was carried out according to the manufacturer’s recommendations.

### Nanopore Sequencing

Nanopore Sequencing was carried out on a MinION sequencer using FLO-FLG001 flongles or Minion R9.4 flow cells from ONT. The experiment was set up and run with live base calling performed using MinKNOW software with default configurations.

### Lambda data visualization and analysis

The FASTQ files generated after the completion of the reads were aligned to respective references using Nanopore’s MinKNOW interface without any filtering. Integrated Genomics Viewer (IGV) was used to visualize the reads aligned to reference with default parameters (27). These alignments were manually analyzed to measure the performance of the linked-pair strategy.

### gRNA design for targeted sequencing of specific regions in the human genome

Our method designs the sgRNAs for a list of target regions that we want to cover (e.g., a list of genes) jointly (Supplementary Figure 2). For each target region, we cover it using *n* sub-regions (*s_1_*, *s_2_*, …, *s_n_*) of fixed lengths, where *s*_1_ starts before the target region, *s_n_* starts after the region, and each region has a length that can be covered by a single sequencing read (Supplementary Figure 1A). Within each sub-region, we define a linker-search area (LSA), which is where we search for a suitable location for a linker (Supplementary Figure 2A). The size of the LSAs is a parameter for trading off between high flexibility (with larger LSAs) and low searching time (with smaller LSAs).

Inside each LSA, we first enumerate all 20-mers (i.e., length-20 sequences) that are followed by the PAM (NGG) contained in the LSA as a candidate gRNA target site (Supplementary Figure 2B). These candidates are then subjected to several rounds of filtering (Supplementary Figure 2B). First, to achieve good enrichment of sequencing reads at our target regions, we considered only candidate gRNA target sites with a unique sequence in the reference genome. Second, since gRNA targeting is less effective for sites with extreme GC content, we further remove 20-mers with a GC content lower than 40% or higher than 60%. Third, we estimate the off-target effects of each gRNA by computing two measures, h1, and h2, defined respectively as the number of 20mer NGGs in the genome sharing 12mer NGG with the candidate gRNA and the number of 20mer NGGs in the genome sharing 8mer NGG with the candidate gRNA but not sharing 12mer NGG. Then we compute the off-target score as w1 * h1 + w2 * h2, where w1 and w2 are weights given to the two cases respectively. In our experiments, w1 and w2 were set to 256/257 and 1/257, respectively. We remove candidate gRNAs with an off-target score of more than 20. Finally, to avoid ineffective gRNA targeting due to genetic variants, we remove all gRNA candidates that overlap known genetic variants.

From the remaining gRNA candidates, we look for pairs of them that can generate linkers (Supplementary Figure 2C). Each pair of candidate gRNAs on opposite DNA strands in an LSA is considered a valid pair if the resulting linker i) has a length of 300-500bp, to match the expected products of our experimental protocol, ii) is unique in the whole genome, enabling unambiguous *de novo* sequence assembly, and iii) contains at least one heterozygous SNV or indel, to enable haplotype reconstruction.

All the linkers from the valid gRNA pairs are then summarized in a directed acyclic graph, where each node is a linker (Supplementary Figure 2D). A linker node X has a directed edge to linker node Y if the starting coordinate of Y is larger than the ending coordinate of X by 10-20 kbp, such that the corresponding DNA fragment can be covered by a single sequencing read. With this graph constructed, the goal is then to find a path that starts from a linker node within *s*_1_ and ends at a linker node within *s_n_*. In our general algorithm (that allows gRNAs that are not unique in the genome), exactly which path and the corresponding set of gRNAs to take for each target region is decided by considering the graphs of all the target regions jointly, which encourages the reuse of gRNAs to cover multiple target regions, and generation of DNA fragments that are close to target read length (15 kbp).

When we applied our method to design gRNAs for sequencing the selected cancer-related genes in NA12878, we partitioned genes into sub-regions of the same length as the target fragment length for sequencing on the Oxford Nanopore platform, namely 15 kbp. As a result, the number of sub-regions for a gene of length is ⌈*L*/15000⌉ + 2. The LSAs were also set to 15 kbp to search for linkers inside the whole sub-regions. We used variant calls with phase-resolved haplotype information for the individual NA12878 from Genome in a Bottle (GIAB) (v.3.3.2) (https://ftp-trace.ncbi.nlm.nih.gov/ReferenceSamples/giab/release/NA12878_HG001/latest/) (18). To further avoid interference with large structure variants, we checked large SVs reported in existing studies to ensure our gRNAs do not overlap them (28–32).

### Generation of human linked-pair fragments

Each 900 ng of high molecular weight NA12878 DNA was preprocessed by first blocking at the 3’ ends with Klenow (exo-) (NEB) in the presence of 10 µM dideoxynucleotides and 1X NEBuffer 3.1 for 30m at 37°C. Next, 3U of rSAP (NEB) was added to deplete the nucleotides and dephosphorylate the 5’ ends and incubated at 37°C for 30m followed by a deactivation step at 65°C for 15m.

Linked pair fragments for our design were generated by sequential fragmentation of short genes followed by long genes. First, the cas9-sgRNA complex targeting all short genes was formed by incubating 37.5 pmol sgRNA with 400ng Cas9 Nuclease (NEB) at 37°C for 15m. This complex was then added to the pre-processed DNA, mixed well and the cleaving reaction was carried out at 1X NEBuffer 3.1 for an hour at 37°C followed by a deactivation step at 65°C for 10 m.

In a separate tube, 400ng Cas9 H840A (IDT) was incubated with 37.5pmol sgRNA for long genes and at 37°C for 15m. This complex was then added to the cleaved DNA from the earlier step and a nicking reaction was carried out at 37°C for 2h at 1X NEBuffer 3.1. An extension step followed where the nicked DNA was incubated with 200uM dNTPs and 3U of Vent (exo-) at 72°C for 1 hour. Linked-pair fragments were purified twice with AMPURE XP beads before sequencing library preparation.

### Human data analysis

Adapters were trimmed from the raw sequencing reads using Porechop (v0.2.4) (https://github.com/rrwick/Porechop) with default parameter settings except for skipping splitting reads based on middle adapters (“--no_split”) since the library preparation did not use middle adapters. Porechop can automatically infer the adapters by searching from pre-built adapter sets. The adapter-trimmed reads were aligned to GRCh38 using Minimap2 (v2.17) (33) with default parameter settings. Duplicate reads and secondary and supplementary alignments were excluded. By default, we used all aligned reads without filtering in our analyses. Gene annotations were obtained from NCBI RefGene release 109. The alignments were visualized using IGV tools (27).

### Human reference-based haplotype phasing

We performed the reference-based haplotype phasing of the genetic variants within the linkers only and within the full gene length regions, respectively. All our haplotype phasing results were compared to the GIAB haplotypes.

To phase the haplotypes of the variants within the linkers, we started with calling variants as follows. Using the read alignments, at each site in the reference, we computed the frequency of observing each of the four nucleotides, a deletion, or an insertion on the aligned reads. For called bases, only those with a quality >= 10 were included. For each site, if the total number of aligned reads is *n* and the number of reads that support the two most frequent alleles are *n*_1_ and *n*_2_, respectively, we called the site a heterozygous variant if (i) *n*_1_ + *n*_2_ ³ 10, (ii) (*n*_1_ + *n*_2_) ³ *n*/2, and (iii) *n*_1_£ 2*n*_2_ and *n*_2_ £ 2*n*_1_. After identifying all the heterozygous variants, we assigned reads to the alleles that they contained to identify alleles from different loci that co-occurred in the reads.

Next, we created a haplotype graph based on the heterozygous variants. For the *i*-th variant *v*_*i*_, we denote the two most frequent alleles as 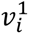 and 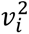. Each of these alleles was represented by a node in the graph. For any two alleles 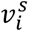 and 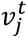, if they co-occur in at least one sequencing read, an edge was drawn between them and the edge received a weight 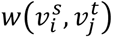 equal to the number of sequenced reads that they co-occur in. Next, we analyzed the edge weights to identify the significant ones as follows. For any two variant sites *v*_*i*_ and *v*_*j*_ in which the edges between them had a total weight of at least 10, we built a 2×2 contingency table 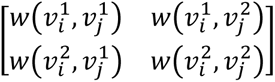 and performed a two-sided Fisher’s exact test based on it. If the p-value was smaller than 0.05, we considered the alleles at the two variant sites significantly correlated and assigned 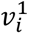 and 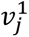 to the same haplotype and 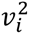 and 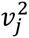 to the same haplotype if 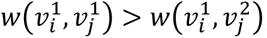; otherwise we assigned 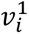 and 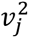 to the same haplotype and 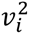 and 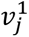 to the same haplotype. After that, we performed another round of testing to improve the accuracy of the inferred haplotypes. For each variant site *v*_*i*_, we collected all the computed p-values involving this site and corrected these p-values using the Bonferroni method. If less than one-third of the resulting corrected p-values were smaller than 0.05, we excluded the site from the inferred haplotypes.

To phase the haplotypes of the variants within the whole gene body regions, we identified heterozygous SNVs anywhere in the gene body regions based on the alignments of the sequencing reads to the reference. Since base calling and sequence alignment at homopolymeric and highly repetitive regions are less reliable, we filtered out variants within regions that contained at least five copies of consecutive 1 bp or 2 bp repeats.

### Human *de novo* haplotype phasing

We designed a grouping and consensus-based method for *de novo* haplotype phasing. Sequencing reads were first grouped together based on the occurrence of the designed gRNAs on their expected linker regions. Specifically, within the first and last 500 bp of each read, we searched for matches of gRNAs (including PAM) with an editing distance of no more than 4. Both the sequences and their reverse complements were considered. For each pair of gRNAs (g1, g2) defining a linker region, we collected reads with matches to either g1 or g2. Based on the matching results, we extracted the parts of the reads potentially overlapping each linker. After this grouping, we performed a multiple sequence alignment for each group of read parts using MAFFT (v7.490) (34), the results of which were used to generate a consensus sequence for each linker region by majority voting. To avoid unreliable results at positions with a small number of supporting reads, we excluded positions with fewer than 30 reads having the two most frequent alleles in total. After getting the consensus sequences of the linkers, the reads were aligned to them to construct haplotype graphs similar to the reference-based method.

## DATA AVAILABILITY

The raw data in this report has been deposited to Sequence Reads Archive and is available at BioProjectID: PRJNA1017960. The source codes for gRNA design, Linked-seq data analysis, and haplotype phasing are available in the GitHub repository https://github.com/Yip-Lab/Linked-seq.git.

## SUPPLEMENTARY DATA

Supplementary Data are available online.

## REFERENCES

1. Wenger, A.M., Peluso, P., Rowell, W.J., Chang, P.-C., Hall, R.J., Concepcion, G.T., Ebler, J., Fungtammasan, A., Kolesnikov, A. and Olson, N.D. (2019) Accurate circular consensus long-read sequencing improves variant detection and assembly of a human genome. Nature biotechnology, 37, 1155–1162.

2. Wang, Y., Zhao, Y., Bollas, A., Wang, Y. and Au, K.F. (2021) Nanopore sequencing technology, bioinformatics and applications. Nature Biotechnology, 39, 1348–1365.

3. (2023) Method of the Year 2022: long-read sequencing. Nature Methods, 20, 1–1.

4. Chen, Y., Nie, F., Xie, S.-Q., Zheng, Y.-F., Dai, Q., Bray, T., Wang, Y.-X., Xing, J.-F., Huang, Z.-J. and Wang, D.-P. (2021) Efficient assembly of nanopore reads via highly accurate and intact error correction. Nature Communications, 12, 60.

5. Rang, F.J., Kloosterman, W.P. and de Ridder, J. (2018) From squiggle to basepair: computational approaches for improving nanopore sequencing read accuracy. Genome biology, 19, 90.

6. Miga, K.H., Koren, S., Rhie, A., Vollger, M.R., Gershman, A., Bzikadze, A., Brooks, S., Howe, E., Porubsky, D., Logsdon, G.A. et al. (2020) Telomere-to-telomere assembly of a complete human X chromosome. Nature, 585, 79–84.

7. Chaisson, M.J., Sanders, A.D., Zhao, X., Malhotra, A., Porubsky, D., Rausch, T., Gardner, E.J., Rodriguez, O.L., Guo, L. and Collins, R.L. (2019) Multi-platform discovery of haplotype-resolved structural variation in human genomes. Nature communications, 10, 1784.

8. McFarland, K.N., Liu, J., Landrian, I., Godiska, R., Shanker, S., Yu, F., Farmerie, W.G. and Ashizawa, T. (2015) SMRT sequencing of long tandem nucleotide repeats in SCA10 reveals unique insight of repeat expansion structure. PloS one, 10, e0135906.

9. Wenzel, A., Altmueller, J., Ekici, A.B., Popp, B., Stueber, K., Thiele, H., Pannes, A., Staubach, S., Salido, E. and Nuernberg, P. (2018) Single molecule real time sequencing in ADTKD-MUC1 allows complete assembly of the VNTR and exact positioning of causative mutations. Scientific reports, 8, 4170.

10. Payne, A., Holmes, N., Clarke, T., Munro, R., Debebe, B.J. and Loose, M. (2021) Readfish enables targeted nanopore sequencing of gigabase-sized genomes. Nature biotechnology, 39, 442–450.

11. Höijer, I., Tsai, Y.C., Clark, T.A., Kotturi, P., Dahl, N., Stattin, E.L., Bondeson, M.L., Feuk, L., Gyllensten, U. and Ameur, A. (2018) Detailed analysis of HTT repeat elements in human blood using targeted amplification-free long-read sequencing. Human Mutation, 39, 1262–1272.

12. Schultzhaus, Z., Wang, Z. and Stenger, D. (2021) CRISPR-based enrichment strategies for targeted sequencing. Biotechnology Advances, 46, 107672.

13. Gabrieli, T., Sharim, H., Fridman, D., Arbib, N., Michaeli, Y. and Ebenstein, Y. (2018) Selective nanopore sequencing of human BRCA1 by Cas9-assisted targeting of chromosome segments (CATCH). Nucleic acids research, 46, e87–e87.

14. Quan, J., Langelier, C., Kuchta, A., Batson, J., Teyssier, N., Lyden, A., Caldera, S., McGeever, A., Dimitrov, B. and King, R. (2019) FLASH: a next-generation CRISPR diagnostic for multiplexed detection of antimicrobial resistance sequences. Nucleic acids research, 47, e83–e83.

15. Gilpatrick, T., Lee, I., Graham, J.E., Raimondeau, E., Bowen, R., Heron, A., Downs, B., Sukumar, S., Sedlazeck, F.J. and Timp, W. (2020) Targeted nanopore sequencing with Cas9-guided adapter ligation. Nature biotechnology, 38, 433–438.

16. Kline, C.N., Joseph, N.M., Grenert, J.P., van Ziffle, J., Talevich, E., Onodera, C., Aboian, M., Cha, S., Raleigh, D.R. and Braunstein, S. (2017) Targeted next-generation sequencing of pediatric neuro-oncology patients improves diagnosis, identifies pathogenic germline mutations, and directs targeted therapy. Neuro-oncology, 19, 699–709.

17. Aaltonen, L.A., Abascal, F., Abeshouse, A., Aburatani, H., Adams, D.J., Agrawal, N., Ahn, K.S., Ahn, S.-M., Aikata, H., Akbani, R. et al. (2020) Pan-cancer analysis of whole genomes. Nature, 578, 82–93.

18. Zook, J.M., Chapman, B., Wang, J., Mittelman, D., Hofmann, O., Hide, W. and Salit, M. (2014) Integrating human sequence data sets provides a resource of benchmark SNP and indel genotype calls. Nature biotechnology, 32, 246–251.

19. Ionita-Laza, I., McCallum, K., Xu, B. and Buxbaum, J.D. (2016) A spectral approach integrating functional genomic annotations for coding and noncoding variants. Nature genetics, 48, 214–220.

20. Wick, R.R., Judd, L.M., Gorrie, C.L. and Holt, K.E. (2017) Completing bacterial genome assemblies with multiplex MinION sequencing. Microbial genomics, 3.

21. Slizovskiy, I.B., Oliva, M., Settle, J.K., Zyskina, L.V., Prosperi, M., Boucher, C. and Noyes, N.R. (2022) Target-enriched long-read sequencing (TELSeq) contextualizes antimicrobial resistance genes in metagenomes. Microbiome, 10, 185.

22. Koboldt, D.C. (2020) Best practices for variant calling in clinical sequencing. Genome Medicine, 12, 91.

23. Abid, H.Z., Young, E., McCaffrey, J., Raseley, K., Varapula, D., Wang, H.-Y., Piazza, D., Mell, J. and Xiao, M. (2021) Customized optical mapping by CRISPR–Cas9 mediated DNA labeling with multiple sgRNAs. Nucleic Acids Research, 49, e8–e8.

24. McCaffrey, J., Young, E., Lassahn, K., Sibert, J., Pastor, S., Riethman, H. and Xiao, M. (2017) High-throughput single-molecule telomere characterization. Genome research, 27, 1904–1915.

25. Shalem, O., Sanjana, N.E., Hartenian, E., Shi, X., Scott, D.A., Mikkelsen, T.S., Heckl, D., Ebert, B.L., Root, D.E. and Doench, J.G. (2014) Genome-scale CRISPR-Cas9 knockout screening in human cells. Science, 343, 84–87.

26. Gilpatrick, T., Wang, J.Z., Weiss, D., Norris, A.L., Eshleman, J. and Timp, W. (2023) IVT generation of guideRNAs for Cas9-enrichment nanopore sequencing. bioRxiv.

27. Robinson, J.T., Thorvaldsdóttir, H., Winckler, W., Guttman, M., Lander, E.S., Getz, G. and Mesirov, J.P. (2011) Integrative genomics viewer. Nature biotechnology, 29, 24–26.

28. Sudmant, P.H., Rausch, T., Gardner, E.J., Handsaker, R.E., Abyzov, A., Huddleston, J., Zhang, Y., Ye, K., Jun, G. and Hsi-Yang Fritz, M. (2015) An integrated map of structural variation in 2,504 human genomes. Nature, 526, 75–81.

29. Li, L., Leung, A.K.-Y., Kwok, T.-P., Lai, Y.Y., Pang, I.K., Chung, G.T.-Y., Mak, A.C., Poon, A., Chu, C. and Li, M. (2017) OMSV enables accurate and comprehensive identification of large structural variations from nanochannel-based single-molecule optical maps. Genome biology, 18, 1–19.

30. Fan, X., Chaisson, M., Nakhleh, L. and Chen, K. (2017) HySA: a Hybrid Structural variant Assembly approach using next-generation and single-molecule sequencing technologies. Genome research, 27, 793–800.

31. Kidd, J.M., Cooper, G.M., Donahue, W.F., Hayden, H.S., Sampas, N., Graves, T., Hansen, N., Teague, B., Alkan, C. and Antonacci, F. (2008) Mapping and sequencing of structural variation from eight human genomes. Nature, 453, 56–64.

32. Pendleton, M., Sebra, R., Pang, A.W.C., Ummat, A., Franzen, O., Rausch, T., Stütz, A.M., Stedman, W., Anantharaman, T. and Hastie, A. (2015) Assembly and diploid architecture of an individual human genome via single-molecule technologies. Nature methods, 12, 780–786.

33. Li, H. (2018) Minimap2: pairwise alignment for nucleotide sequences. Bioinformatics, 34, 3094–3100.

34. Nakamura, T., Yamada, K.D., Tomii, K. and Katoh, K. (2018) Parallelization of MAFFT for large-scale multiple sequence alignments. Bioinformatics, 34, 2490–2492.

